# Generalisation behaviour of predators toward warning signals displayed by harmful prey: answers from a videogame played by humans

**DOI:** 10.1101/409557

**Authors:** Mónica Arias, David Griffiths, Mathieu Joron, John Davey, Simon Martin, Chris Jiggins, Nicola Nadeau, Violaine Llaurens

## Abstract

The persistence of several warning signals in sympatry is a puzzling evolutionary question because selection favours convergence of colour patterns among toxic species. Such convergence is shaped by predators’ reaction to similar but not identical stimulus, *i.e.* generalisation behaviour. However, studying generalisation behaviour in complex natural communities of predators is challenging, and is thus generally limited to simple variations of prey colour patterns. Here, we used humans as surrogate predators to investigate generalisation behaviours on two prey communities with different level of warning signals complexity. Humans’ generalisation capacities were estimated using a computer game simulating a simple (4 morphs) and a complex (10 morphs) community of defended (associated with a penalty) and palatable butterflies. Colour patterns used in the game are actually observed in natural populations of the defended butterflies *H. numata*, and generalisation behaviour of natural predator’s communities on these colour patterns have previously been investigated in the wild, allowing direct comparison with human behaviour. We investigated human predation behaviour by recording attack rates on the different defended and palatable colour patterns, as well as player survival time (i.e. score). Phenotypic similarity among the different colour patterns was precisely quantified using a custom algorithm accounting for both colour and pattern variations (CPM method). By analysing attack behaviours of 491 game players, we found that learning was more efficient in the simple prey community. Additionally, profitable prey gained protection from sharing key visual features with unprofitable prey in both communities while learning, in accordance with natural predator behaviours. Moreover, other behaviours observed in natural predators, such as colour neophobia, were detected in humans and shaped morph vulnerability during the game. Similarities between our results in humans and the reaction of natural predator communities to the same colour patterns validate our video-game as a useful proxy to study predator behaviour. This experimental set-up can thus be compared to natural systems, enabling further investigations of generalisation on mimicry evolution.

## Introduction

Predation exerts a strong selective pressure on prey appearance, promoting for example the evolution of colour patterns that match their background (*i.e.* crypsis), or of warning signals advertising toxicity. Such warning signals are often conspicuous colour patterns, easy to distinguish from their background (Cott, 1940; Poulton, 1890). Despite the influence of innate responses (Rubinoff & Kropach, 1970; Smith, 1977), most predators need to sample aposematic prey several times before learning the association between appearance and distastefulness (Alatalo & Mappes, 1996; Gittleman & Harvey, 1980; Lindstrom, Alatalo, Mappes, Riipi, & Vertainen, 1999; Sillén-Tullberg, 1985). Therefore, the larger number of toxic prey displaying a common warning signal, the smaller is the individual predation risk (Müller, 1878). Such positive frequency selection on conspicuous signals often promotes evolutionary convergence between distantly-related toxic species living in sympatry, thus forming mimicry rings of species sharing similar appearance (Lindström, Alatalo, Lyytinen, & Mappes, 2001; Mallet & Gilbert, 1995; Rowland, Ihalainen, Lindstrom, Mappes, & Speed, 2007). Visual signals can vary among co-mimics, as shown in Arctiid moth (Ojala, Lindström, & Mappes, 2007) and Dendrobatidae frogs (Rojas & Endler, 2013). Predator reaction to these phenotypic differences (*i.e.* generalisation) then shapes the evolution of convergent warning signals. Generalisation capacities can range from very narrow, imposing a strong purifying selection on resemblance, to a broader spectrum tolerating some deviations from the initially learned signal (Ruxton, Franks, Balogh, & Leimar, 2008). Narrow generalisation capacities favouring evolution of high resemblance among co-mimics (Rowland, et al., 2010) can be exerted by experienced (Rowe, Lindström, & Lyytinen, 2004) and bold predators (Thomas, Marples, Cuthill, Takahashi, & Gibson, 2003) and can be triggered by scarce alternative palatable prey (Rowe, et al., 2004). Generalisation is thought to be broad when co-mimics exhibit high levels of distastefulness (Duncan & Sheppard, 1965) and when imperfect mimics are relatively rare or are not highly profitable (Penney, Hassall, Skevington, Abbott, & Sherratt, 2012; Sherratt, 2002). Thus, generalisation is not only determined by cognitive capacities and sensitivities of the predator, but it is also shaped by prey community composition. In complex communities where multiple distinct aposematic signals co-exist, predator learning is slower and generalisation might be less strict, conferring similar protection to various prey displaying some common visual elements in their colour pattern (Ihalainen, Rowland, Speed, Ruxton, & Mappes, 2012). Predators may indeed categorise potential prey as profitable or not according to few key visual elements shared by defended species (Beatty et al., 2004; Chittka & Osorio, 2007). In the experiments mentioned above, generalisation has mainly been tested using artificially modified colour patterns, instead of natural prey patterns that predators may encounter in wild communities. Some field studies have used patterns displayed by natural prey, but that do not co-exist in the same locality (Amézquita, Castro, Arias, González, & Esquivel, 2013), therefore, natural predators do not learn those specific combinations simultaneously.

Here, we focus on the unpalatable butterfly *Heliconius numata* that exhibits an exceptional polymorphism in wing colour pattern. Within population, up to five discrete colour patterns can be maintained, each belonging to a different mimicry ring (Brown & Benson, 1974). *H. numata* butterflies generally display tiger colour patterns with orange, black and sometimes yellow. Between these different morphs, those displaying yellow patches might be more conspicuous given the high contrast between yellow and black (Llaurens, Joron, & Théry, 2014). Conspicuousness is indeed supposed to be enhanced by highly contrasting brightness between colour patches and background (Endler, 1992). Using artificial butterflies displaying actual patterns and placed in natural populations, Arias et al. (2016) investigated the effect of phenotypic similarity on generalisation behaviour by measuring attack rate on artificial butterflies displaying local mimetic *H. numata* colour patterns, as well as intermediate patterns obtained from crosses between those local mimetic patterns. Attack rates on intermediate morphs were lower when (1) resemblance to a locally mimetic morph was higher, and when (2) appearance of parental morphs was more similar. Altogether these results highlight the generalisation capacities of predators toward local variations in warning signals. These correlations were stronger when colour perception was included in the estimation of resemblance, stressing the important role of colour in generalisation. However, to investigate large range of phenotypic variations, artificial prey approaches are limited because attack rates are extremely low (5.34% in Arias *et al.* 2016, 12.72% in Chouteau and Angers 2011, 2.3% of avian predators traces in Noonan and Comeault 2009, 11.77% in Finkbeiner *et al* 2012 and 4.03% in Merrill *et al.* 2012). Although using artificial prey in natural populations allowed us to directly estimate selection exerted by natural communities of predators such as different bird species, tests with humans can cover a larger and more detailed range of phenotypic variation and have proven useful in the study of generalisation (Sherratt, Whissell, Webster, & Kikuchi, 2015). For instance, experiments with humans have been used to test the hypothesis that some colour patterns are cryptic at a distance but involve signalling at closer range (*i.e.* distance dependent dual function) (Bohlin et al., 2012), the evolution of non-conspicuous traits signalling unprofitability (Sherratt & Beatty, 2003) and evolution of slow movement behaviours in protected prey (Sherratt, Rashed, & Beatty, 2004). Humans produced similar reaction to birds in several behaviours: for example, great tits as well as humans can use aggregation of prey as a signal of prey unprofitability when generalising (Beatty, Bain, & Sherratt, 2005). Moreover, under the same experimental design, blue tits (Kazemi et al., 2014) and humans (Sherratt et al., 2015) distinguished profitable from unprofitable prey similarly, focusing more on colour cues that seem more striking (salient), than prey pattern or shape information. These studies show that human generalisation resembles natural predator behaviour and can allow us to tackle specific questions otherwise difficult to achieve in other experimental conditions.

Here, we tested generalisation behaviour towards aposematic colour patterns displayed by *H. numata*, using a computer game played by humans. We first aim at comparing human to natural predator community generalisation behaviour and then at investigating generalisation behaviour in communities with different levels of waring signal diversity using the video-game as a proxy of predator behaviour. We used two versions of the game: (1) using the ten morphs tested in the field experiment described in Arias et al. (2016) (referred to as complex community); and (2) using only four out of those morphs (simple community). Moreover, humans can be considered as naïve predators, so that our experimental set up allows us to investigate the effect of phenotypic similarity during and after learning.

In the game, players had the role of hungry predators that need to quickly eat as many profitable butterflies as possible to survive. In each trial, two butterfly morphs were randomly chosen as “defended morphs” and associated with a penalty in survival time. We recorded attack number on each palatable and defended morphs as well as the time that the player stayed alive in the game (player score). We estimated phenotypic similarity among butterfly colour patterns using a custom algorithm accounting for both colour and pattern variations (i.e. CPM method, see Arias et al. 2016). We then compared human and natural predator responses by testing 1) whether protection of a profitable prey was enhanced by phenotypic similarity with defended prey and 2) whether the phenotypic proximity within each pair of defended morphs enhanced their protection. We then investigated 3) learning behaviour of players, and 4) the effect of warning signal diversity on such learning process.

## Materials and methods

### Butterfly images

To investigate the generalisation capacities of human observers, we used (1) a complex community including 5 intermediate (heterozygous) phenotypes and 5 corresponding mimetic homozygous phenotypes including a large part of phenotypic variation observed within the polymorphic species *H. numata* (see Arias et al. (2016) for more detail). Additionally, we used a subset of 4 of those mimetic homozygous phenotypes (namely *tarapotensis, bicoloratus, aurora* and *arcuella*) to build (2) a simple warning signal community. Three of these homozygous phenotypes are actually sympatric in natural populations. These butterflies were photographed under standard light conditions.

### Computer game

The computer game *Hungry birds v2* was developed from a previous version designed for evolution outreach (*Hungry birds v1* was display on the *Heliconius* stand of the Royal Society Exhibition 2014 in London and is still available from the *Heliconius* community website, http://www.heliconius.org/summer-science/evolving-butterflies-game/). Players were asked to catch butterflies by touching them on the screen, simulating hungry predators from the tropical forest. In each trial, two morphs were randomly assigned as defended. Once a defended morph was touched by a player, a warning message was displayed on the screen stating ‘Ugh! That butterfly tasted disgusting’. Players were then prevented from eating any more butterflies within the next 2 seconds as penalty. At the top of the screen, players could see their constantly decreasing life bar. It increased after catching a profitable butterfly (life gain), and quickly decreased after touching a defended butterfly (cost in survival associated with attack of a defended butterfly). To mimic natural conditions, a maximum of five butterflies appeared simultaneously in the screen, limiting direct comparison between morphs. Player’s motivation stem from preventing life bar to get too low (mimicking hunger level) and getting a high score (associated to the time each player stayed alive in the game).

### Volunteer players

In June 2015 and March 2017, we invited the general public visiting the Evolution Gallery (Grande Galerie de l’Evolution) at the National Museum of Natural History in Paris (France), to play the game. *Hungry Birds v2* was loaded on a Raspberry *Pi*, and was accessed by a tablet through Wi-Fi. We invited people from all ages and we tried to sample both sexes evenly. First, a short explanation of the rules of the game was given. Once the game was finished, explanations about the aim of the experiment and the mimetic interactions occurring among *Heliconius* and other colourful Neotropical butterflies were provided and illustrated by actual butterflies and predators displayed at the museum. Players were invited to play two or more times, taking the first time as a familiarization experience. Only players’ age (recorded by class: younger than 10, 10 to 15, 16 to 35, 36 to 50 and older than 50) and number of trials played were recorded to correct for potential bias. The game was thus entirely anonymous and based on volunteering, and no information on the participants was kept except age, following ethical recommendations.

### Estimation of phenotypic distances and rates of attack

Phenotypic distances among morphs were computed following Arias *et. al* (2016), using the Colour Pattern Modelling (CPM) method described in Le Poul *et al.* (2014) and implemented in Matlab (MATLAB, 2012). In CPM, pictures of actual butterflies’ wings were aligned (using rotation, translation and rescaling) to a colour pattern model built recursively, minimizing differences between each colour pattern to the model wing. After alignment, the position of each pixel of the wing image was considered homologous among all individuals. Phenotypic variations were then described by Principal Component Analysis (PCA), using binary values for presence/absence of each of the four colour classes (black, orange, yellow, white) as values for each pixel of the wing image (referred to as binary PCA hereafter). Alternatively, we also performed a “perceptual” PCA, taking into account human perception of butterflies’ colours. Perceptual PCA variables were the relative amount of light captured by each photoreceptor when observing a given colour (*i.e.* quantum catch (Iriel & Lagorio, 2010)), estimated for humans with sensitivity ranging from 400-700 nm (Dartnall, Bowmaker, & Mollon, 1983). To extract the quantum catches, we applied the method described in Vorobyev and Osorio (1998), including human colour vision (Dartnall et al., 1983), assuming a Weber fraction of 0.05, human cone photoreceptor ratios (SWS = 1, MWS = 16, LWS = 32 (Dartnall et al., 1983)) and using the reflectance spectra measured on black, orange and yellow patches of actual wings of *H. numata tarapotensis* morph with an AvaSpec-3468 spectrophotometer (Avantes, Apeldoorn, the Netherlands) and a deuterium–halogen light source (DH-2000, Avantes) connected to a 1.5-mm-diameter sensor (FCR-7UV200-2-1.5 × 100, Avantes) inserted in a miniature black chamber. These calculations were performed with the software AVICOL (Gomez, 2006) assuming “small gap” light condition (Théry, Pincebourde, & Feer, 2008).

Müllerian mimicry among toxic species is promoted by the advantage gained from sharing a common warning signal. However this advantage depends on the level of resemblance between co-mimics and on the perception of this resemblance by predators. Therefore, we estimated phenotypic similarities between all morph pairs by computing Euclidian distance between morphs on the 15 PCA components for both binary and perceptual PCA. In the game, the resemblance between the two defended colour patterns differs among trials, because the two defended morphs were randomly chosen among the four (in the simple community) or ten (in the complex community) possible morphs for each trial. This allowed us to test whether defended butterflies benefit from greater protection when they display a more similar colour pattern, because of positive number-dependent selection. We thus computed the phenotypic distance 1) between the two defended morphs (*D*_*def*_) in the trial, and 2) between each stamped palatable butterfly and the most similar defended morph (*D*_*pal*_). The most similar morph was identified based on the phenotypic distance computed both from binary and perceptual PCA independently. Then, statistical analyses related to phenotypic distances were performed independently for the results obtained by each PCA calculations.

Player variables (ID number associated to each participant trial, player age and trial score measured as trial duration) and trial variables (ID of the two defended morphs, total number and ID of butterflies consumed, as well as order of stamped butterflies) were recorded. Trials with less than 4 or 10 butterflies stamped (for simple and complex community game respectively) were discarded, in order to analyse only trials where players are likely to have encountered most of the community diversity. First, by pooling trials of all players for each community, we used the butterfly attack order to compute 1) the attack peak (height and position) on defended and palatable colour patterns. We then 2) compared the time taken by players to learn to avoid defended morphs (*i.e.* position of the peak) and 3) the strength of the avoidance after learning (*i.e.* slope after the attack peak, assuming that a steeper decrease indicates a quicker memorization of the warning signal). All these calculations were performed on the density curves of butterfly attack, built from histograms of butterfly attack position. To dissect the importance of previous experience on butterfly consumption, we divided trial each trail in two phases: (1) “hunting while learning” phase (before the peak) and (2) “hunting by experts” phase (after the peak). We then compared defended and undefended butterflies consumption rate while learning and by experts for both communities. Based on the stamped butterflies within each phase of each trial, we calculated for each morph *M*: 1) the general attack proportion: attacks on morph *M* / total number of attacks in the phase and 2) the relative attack rate, similarly as in the field experiment described in Arias et al.(2016): attacks on morph *M* /attacks on the most similarly looking defended morph.

### Statistical analyses

Chi-square tests were applied to compare number of attacks between morphs. Differences between butterflies attacked rate were tested with an ANOVA and a Tukey post-hoc test.

To test whether phenotypic similarity to unprofitable morphs (*D*_*pal*_) conferred any advantage to profitable prey, we fitted linear mixed models (LMM) for simple and complex communities independently, including: participant ID as random factor, relative attack rate on profitable prey as response variable and phenotypic distance and ID of the most similarly looking defended morph, player score and age, presence of yellow in morph and sharing colours with none, one or both defended morphs as explanatory variables. We also tested whether higher similarity between the two unprofitable morphs (*D*_*def*_) increased protection by fitting LMMs assuming: participant ID as random factor, attack rate on defended morphs as response variable and distance between defended morphs, player score and age, total number of attacked butterflies, combination of defended morphs and sharing or not colour between them as explanatory variables. A similar model including defended morph combination instead of phenotypic distance between defended morphs was also fitted. We assumed normal distribution for response variables for GLMs. All statistics were computed using R (R Foundation for Statistical Computing, 2014).

## Results

### Discriminating defended from profitable morphs in contrastingly diverse communities

In 2015, 342 participants played a complex community version of the game (with 10 morphs), out of which 253 players managed to attack at least 10 butterflies (∼74% of participants). In 2017, 149 participants played the simpler community version of the game (with 4 morphs) out of which 145 managed to attack at least 4 butterflies (∼97% of participants).

The attack rate on profitable morphs was significantly higher than on defended ones, in the simple community (2017) (*χ*^2^ =2717, df = 1, *p* < 0.0001) as well as complex community (2015): (*χ*^2^ = 3324.5, df=1, *p* < 0.0001), confirming that players learned to distinguish defended from profitable morphs in both communities.

### Comparing morph conspicuousness

Each morph suffered 8.938 ± 9.97 attacks on average for simple community and 2.883 ± 3.30 attacks for the complex one. The total number of attacks was not significantly different between morphs in either of the communities (simple community: *χ*^2^ = 0, df = 3, *p* = 1; complex community: *χ*^2^ = 0, df = 9, *p* = 1). Defended morphs suffered 0.478 ± 1.13 attacks in complex communities and 1.253 ±1.50 in simple communities. No difference in number of attacks between the two defended morphs was detected in neither of the communities (simple community: *χ*^2^ =4.182, df = 3, *p* = 0.242; complex community: *χ*^2^ = 4.45, df = 9, *p* = 0.879). These results suggest that humans do not exhibit strong avoidance regarding specific colour patterns.

### Learning behaviour

Attack peak for undefended occurred sooner in the simple community (5.3 time units) than complex community (6.45 time units) (fig. 2). In the simple community, player attacks are distributed on fewer morphs, producing more attacks and making the evaluation of all morph profitability occur sooner.

**Figure 1.**
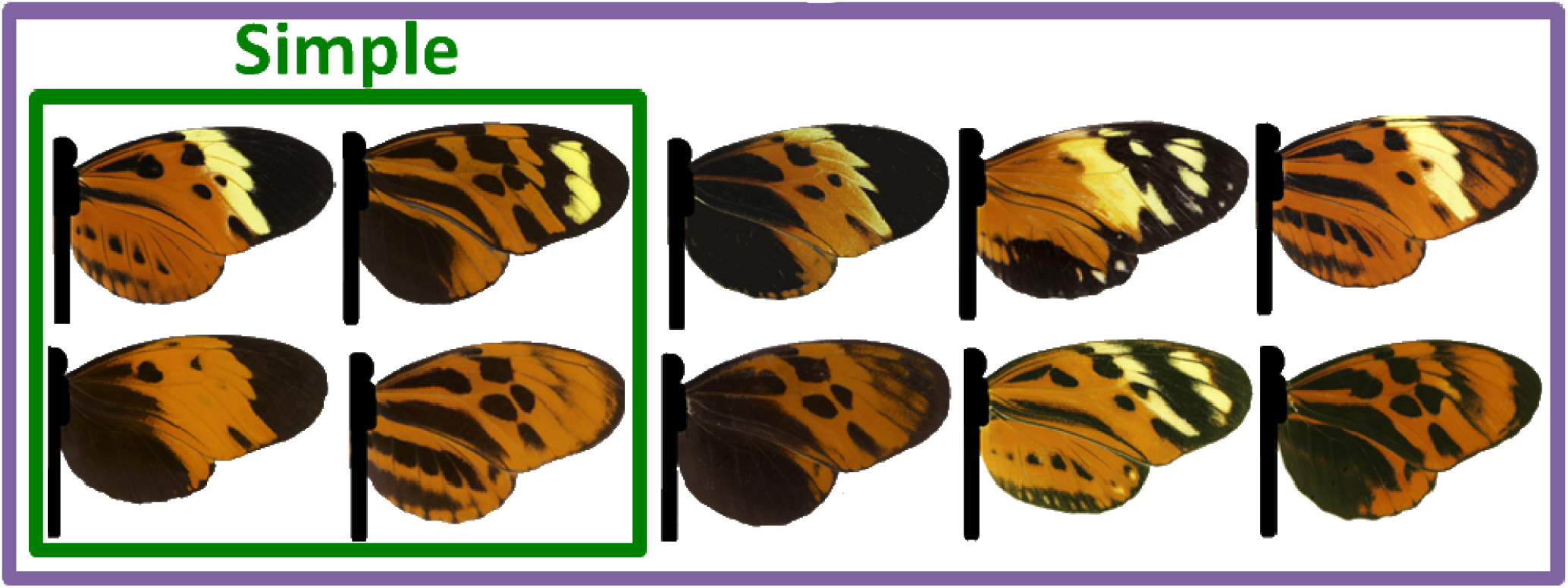
*Heliconius numata* morphs displayed on the simple (green square) and complex (purple square)

**Figure 2.**
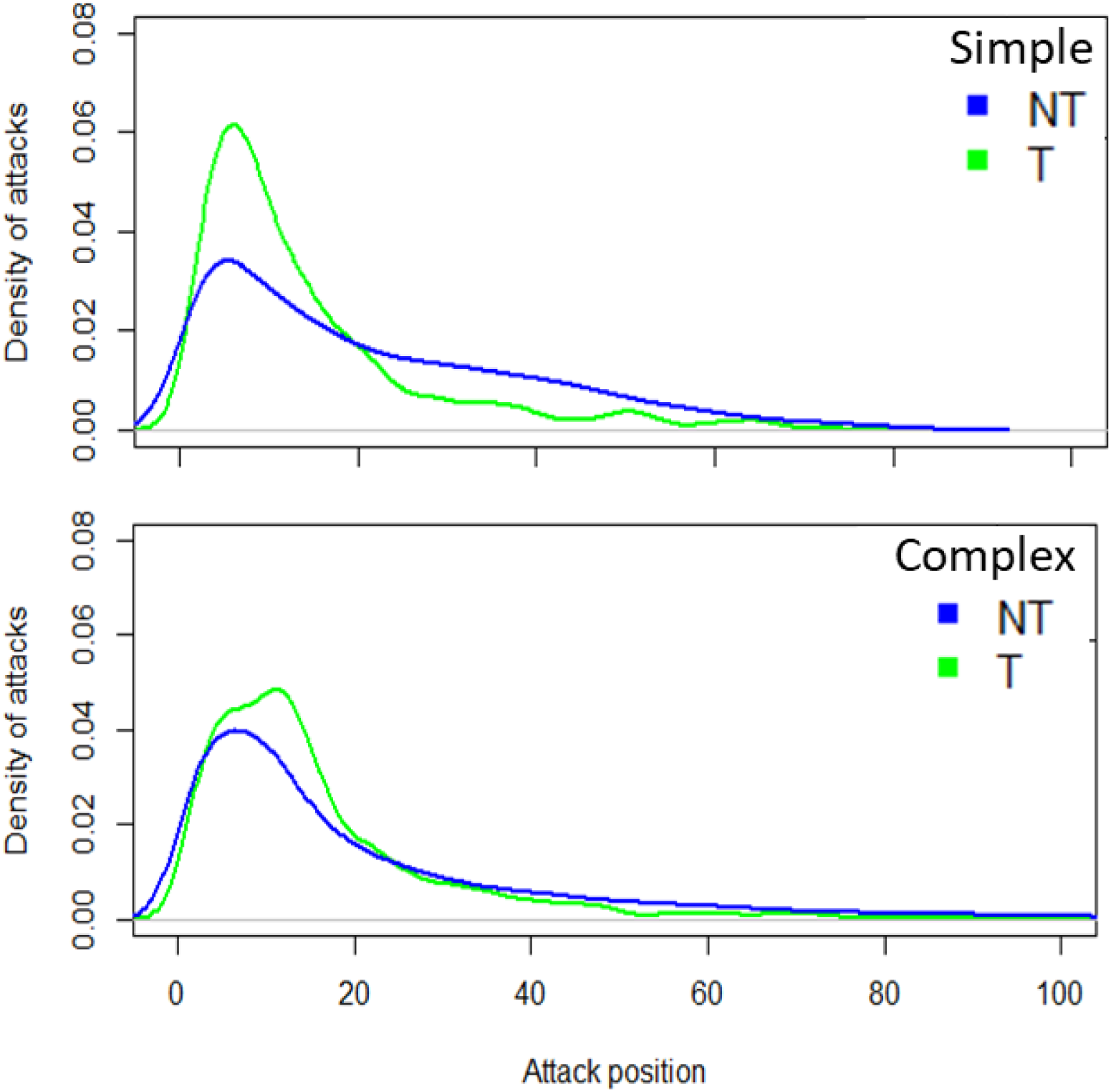
Density of attacks on toxic (T) and non-toxic (NT) morphs along trail duration (plotted as attack position), for simple (top) and complex (bottom) communities.

In the complex community, defended and undefended morphs showed a similar average attack peak height (average peak height for defended: 0.05) (average peak height for undefended: 0.04). However, attacks on defended morphs decreased two times faster than for undefended morphs (Fig. 2; slope: undefended: -0.0017, defended: -0.00353). In the simple community, peak slope was also larger for defended (slope:-0.002) than for undefended morphs (slope: -0.0007). However, peak height was nearly two fold larger for defended than for undefended morphs when less morphs were present in the community (defended: 0.0613; undefended: 0.034).

In both simple and complex communities, humans attacked less defended than undefended butterflies during learning but also as experts (*F* = 125.9, df = 7, *p* < 0.001, Fig. 3). However, the differences between attack rates on defended and palatable prey was larger in the simple community than in the complex community, as suggested by peak height differences (Fig. 2), even after correcting by the proportion of defended forms present in each community (two out of 4 in simple and two out of 10 in complex community, Fig. 3).

**Figure 3.**
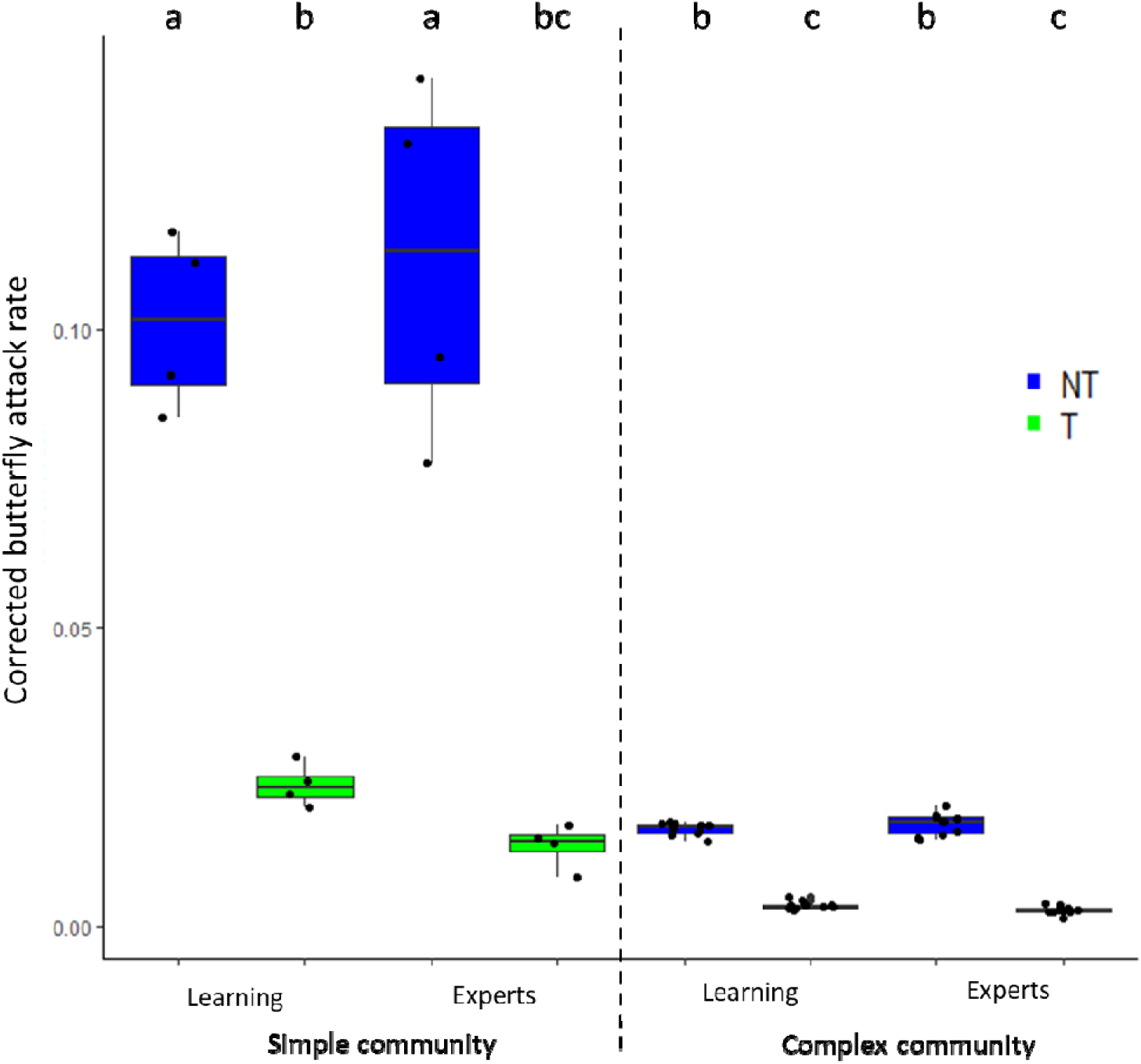
Corrected attack rate on defended (T, in green) and undefended (NT, in blue) butterflies before and after learning in both simple and complex communities. All pairwise comparisons were performed: between phases (learning – experts), defended (T) and undefended (NT) forms for both communities (simple – complex). Letters above boxplots state for difference at α = 0.05.

### Generalisation to defended morph

We tested whether resemblance to the most similar defended morph improved protection by fitting independent linear mixed effect models per distance calculation method, community complexity and learning phase. The best fitting model explaining attack proportion on undefended butterflies when learning on complex community was the only one that included phenotypic distance calculated by binary PCA as an explanatory variable (Table S5). At shorter distance between undefended and defended forms, the proportion of attacks on the undefended morphs was lower. Other explanatory variables included in the best fitting models were: score, positively correlated to the attack proportion in undefended morphs, and included in all the models (Tables S1, S2, S4, S5, S7, S8, S10, S11); closest defended morph ID (relevant after learning in the simple community and for both distance calculation methods, Tables S7 and S8); and the presence of yellow in the undefended form (relevant when learning in the simple community and under both distance calculation methods, Tables S1 and S2).

### Protection gained when toxic morphs looked alike

Expert predators that scored the highest attacked less defended morphs on both simple and complex communities (Tables S9 and S12). Neither phenotypic distance between defended morphs, sharing the presence of yellow by defended morphs, nor specific morphs had any effect on the attack rate suffered by defended forms on neither of the communities (Tables S3, S6, S9, S12).

## Discussion

We used a videogame to investigate generalisation behaviour toward actual warning signals displayed by the polymorphic toxic species *H. numata*. Players, especially those with the highest scores on the game, managed to recognize and avoid wing colour patterns associated with a cost after a given butterfly number sampled, in a similar way that birds can learn to avoid a warning signal associated with a repulsive taste (Pinheiro, 2003; Rowland et al., 2007). Players attacked a similar number of butterflies per morph in both communities. However, participants learned faster to distinguish defended from undefended forms in the simple community (the attack peak for the toxic morphs occurred sooner for the simple community). Moreover, differences in consumption between defended and undefended prey were larger for the simple community. Our results thus suggest that facing a less diverse prey community, favours faster learning by humans. This is consistent with birds response to simple *vs.* complex communities (Ihalainen et al., 2012). Instead, the large diversity of complex communities can hamper predator discrimination capacities, therefore slowing down the learning process.

Participants attacked fewer undefended morphs while learning in the complex community when they were more similarly looking to defended prey. Evidence from field experiments shows how avian predators attacked less morphs when they were more similar to the locally defended morphs (Arias et al., 2016). Our results suggest that when learning to tell apart defended from undefended forms on highly diverse communities, generalisation on common cues could facilitate the task and evolve. Instead, when facing a lower diversity of defended forms, predators can recognise each form independently, without looking for common elements among them.

Smaller distances between defended morphs did not enhance their protection when learning. This contrasts with the Müllerian mimicry expectations, as the similarity among protected prey favours generalisation of warning signal and thus, protection of all prey sharing it (Müller, 1878; Rowland, Hoogesteger, Ruxton, Speed, & Mappes, 2010). This also challenges the importance of distinct cues shared by defended prey in complex communities, where selection for mimicry is strong (Beatty et al., 2004). However, participants may have learnt the specific appearance of all toxic morphs quickly without the need to sample them many times. Therefore, it is possible that the low number of attacks on defended forms makes it difficult to draw conclusions regarding what can enhance their protection before and after learning.

### Variations in human as a relevant proxy for investigating predation behaviours

Interestingly, some players displayed “neophobic” behaviours, *a priori* assuming that unprofitable butterflies must be the brightest or the darkest. This neophobic behaviour can explain why the presence of yellow in undefended morphs, conferred them protection when humans when learning in the simple community. Such innate avoidance of certain colour or shape has been documented in birds (Smith, 1977). Theoretical approaches suggest that such neophobia in predators can benefit rare phenotypes, relaxing negative selection on them and enabling the emergence of a polymorphism in toxic mimetic species (Aubier & Sherratt, 2015; Sherratt, 2011). Human behavioural variation observed in the game could be further investigated using a videogame and an appropriate experimental design. This may lead to relevant evolutionary hypotheses on natural selection acting on warning pattern variations.

### Conclusions

Our videogame played by humans can indeed reproduce natural predator generalisation behaviours when responding to colour pattern variations among the mimetic species *Heliconius numata*. In spite of the different visual systems found in bird predators’ and humans, tools as videogames indeed enable exploring in detail how predators have shaped resemblance between mimics and how other visual protective cues have evolved in wild populations.

## Acknowledgements

The authors would like to thank the sponsors of our Royal Society Summer Science Exhibit in 2014 that made the original development of this game possible. We also thank Fabienne Noe-Stosic for her help in setting up the experiment at the Grande Galerie de l’Evolution of the National Museum of Natural History. We also thank Juan Ulloa, Diego Llusia, Jaqui Ortega and Florence Prunier who helped recruiting volunteers to play. We also thank Martin Stevens, Hannah Rowland, Thomas Lenormand and Marc Théry for comments in previous versions of this manuscript. This work was funded by MA’s PhD grant provided by the Labex BcDIV (LabEx ANR-10-LABX-0003-BCDiv) and by ANR grant DOMEVOL to VL (ANR-13-JSV7-0003-01) and Emergence Program of Paris city council to VL.

